# Craniofacial dysmorphology in Down Syndrome is caused by increased dosage of Dyrk1a and at least three other genes

**DOI:** 10.1101/2022.06.27.497841

**Authors:** Yushi Redhead, Dorota Gibbins, Eva Lana-Elola, Sheona Watson-Scales, Lisa Dobson, Matthias Krause, Karen J. Liu, Elizabeth M. C. Fisher, Jeremy B.A. Green, Victor L.J. Tybulewicz

## Abstract

Down syndrome (DS), trisomy of human chromosome 21 (Hsa21), occurs in 1 in 800 live births and is the most common human aneuploidy. DS results in multiple phenotypes, including craniofacial dysmorphology, characterised by midfacial hypoplasia, brachycephaly and micrognathia. The genetic and developmental causes of this are poorly understood. Using morphometric analysis of the Dp1Tyb mouse model of DS and an associated genetic mouse genetic mapping panel, we demonstrate that four Hsa21-orthologous regions of mouse chromosome 16 contain dosage-sensitive genes that cause the DS craniofacial phenotype, and identify one of these causative genes as *Dyrk1a*. We show that the earliest and most severe defects in Dp1Tyb skulls are in bones of neural crest (NC) origin, and that mineralisation of the Dp1Tyb skull base synchondroses is aberrant. Furthermore, we show that increased dosage of *Dyrk1a* results in decreased NC cell proliferation and a decrease in size and cellularity of the NC-derived frontal bone primordia. Thus, DS craniofacial dysmorphology is caused by increased dosage of *Dyrk1a* and at least three other genes.

**Summary statement:** Craniofacial dysmorphology in mouse models of Down syndrome is caused by increased dosage of at least four genes including *Dyrk1a*, resulting in reduced proliferation of neural crest-derived cranial bone progenitors.

## INTRODUCTION

Down syndrome (DS), trisomy 21, is caused by an additional third copy of human chromosome 21 (Hsa21). DS is the most common human aneuploidy occurring in 1 in 800 live births and results in numerous phenotypes including learning and memory deficits, heart defects and early onset Alzheimer’s disease (Antonarakis, 2017; Bergström et al., 2016; Lott and Dierssen, 2010; Wiseman et al., 2015). Individuals with DS also have a characteristic craniofacial dysmorphology with a near 100% penetrance, characterised by reduction in the dimensions of the midface, with flattened nose bridge (midfacial hypoplasia), shortening of the skull along the anteroposterior axis (brachycephaly), reduction in the dimensions of the lower jaw (micrognathia), altered shape of the skull orbit, and an absence or reduction of permanent teeth (hypodontia) (Antonarakis, 2017; Vicente et al., 2020).

DS is a gene dosage disorder with a third copy of one or more of the ∼230 genes on Hsa21 causing the multiple phenotypes. However, the identity of the causative dosage-sensitive genes is largely unknown and thus the pathological mechanisms underlying DS phenotypes are unclear. Knowledge of such genes and mechanisms is essential for targeted therapies since there are no treatments for most aspect of DS. In particular, the genes and mechanisms driving craniofacial dysmorphology in DS are poorly understood.

The search for dosage-sensitive genes causing DS phenotypes has used both human and mouse genetics. Analysis of rare partial trisomies of Hsa21 has been used to map individual phenotypes to specific regions of the chromosome, initially to the so-called Down syndrome critical region (DSCR) (Delabar et al., 1993; Korenberg et al., 1994; McCormick et al., 1989) but more recently to multiple regions of the chromosome (Korbel et al., 2009; Lyle et al., 2009). Since these partial trisomies are rare, there is insufficient genetic resolution to identify causative genes using this approach. Instead, attention has turned to mouse genetics.

Hsa21 is orthologous to regions on mouse chromosomes 10 (Mmu10), Mmu16 and Mmu17. Using genome engineering, mouse strains have been constructed with duplications of each of these three regions, increasing their copy number from two to three (Lana-Elola et al., 2016; Li et al., 2007; Yu et al., 2010). We generated the Dp1Tyb mouse strain with an extra copy of 23 Mb of Mmu16, the largest of the Hsa21-orthologous regions, containing 142 coding genes, thereby modelling trisomy of around 62% of Hsa21 (Lana-Elola et al., 2016). Analysis of Dp1Tyb mice has shown that they have many DS-like phenotypes, including congenital heart defects, reduced bone density, and deficits in memory, locomotion, hearing and sleep (Chang et al., 2020; Lana-Elola et al., 2021; Lana-Elola et al., 2016; Thomas et al., 2020; Watson-Scales et al., 2018). Notably, we also found that Dp1Tyb mice have craniofacial phenotypes that recapitulate aspects of the human DS dysmorphology, including midfacial hypoplasia, brachycephaly and micrognathia (Toussaint et al., 2021). Similar findings have been reported for the Dp1Yey mouse strain which has an additional copy of the same Mmu16 region (Li et al., 2007; Starbuck et al., 2014).

Here, we use morphometric analysis of a panel of mouse strains, each containing an extra copy of sub-regions of Mmu16, to map the location of genes causing the craniofacial phenotypes in Dp1Tyb mice. We discover that there must be at least four genes whose increased dosage contributes to the craniofacial dysmorphology and show that one of these causative genes is *Dyrk1a*. We demonstrate that the most severe defects in Dp1Tyb skulls are in bones of neural crest (NC) origin and reveal aberrant mineralisation of the Dp1Tyb skull base synchondroses. Furthermore, we show that increased dosage of *Dyrk1a* results in decreased NC cell proliferation and a decrease in the size of the NC-derived frontal bone primordia. Thus, DS craniofacial dysmorphology is caused by increased dosage of *Dyrk1a* and at least three other genes, leading to reduced proliferation of NC cells that generate frontal and facial bones.

## RESULTS

### Genetic mapping identifies four loci causing the craniofacial phenotype in the Dp1Tyb model of DS

To map the location of dosage-sensitive genes that cause the craniofacial dysmorphology of Dp1Tyb mice, we made use of a panel of seven mouse strains that have shorter genomic duplications contained within the Hsa21-orthologous region of Mmu16, thereby breaking up the 23 Mb, 142-gene region duplicated in Dp1Tyb mice into shorter segments (Fig. 1A) (Lana-Elola et al., 2016). We used micro-computed tomography (µCT) to generate 3D images of the skulls of ten 16-week old mice from each of these strains (5 females, 5 males), along with the same number of age- and sex-matched wild-type (WT) control mice. All mice were on a C57BL/6J background. To evaluate the shapes and sizes of the skulls, we carried out landmark-based morphometric analysis on the images, using 68 landmarks for the cranium and 17 landmarks for the mandible.

**Figure 1.**
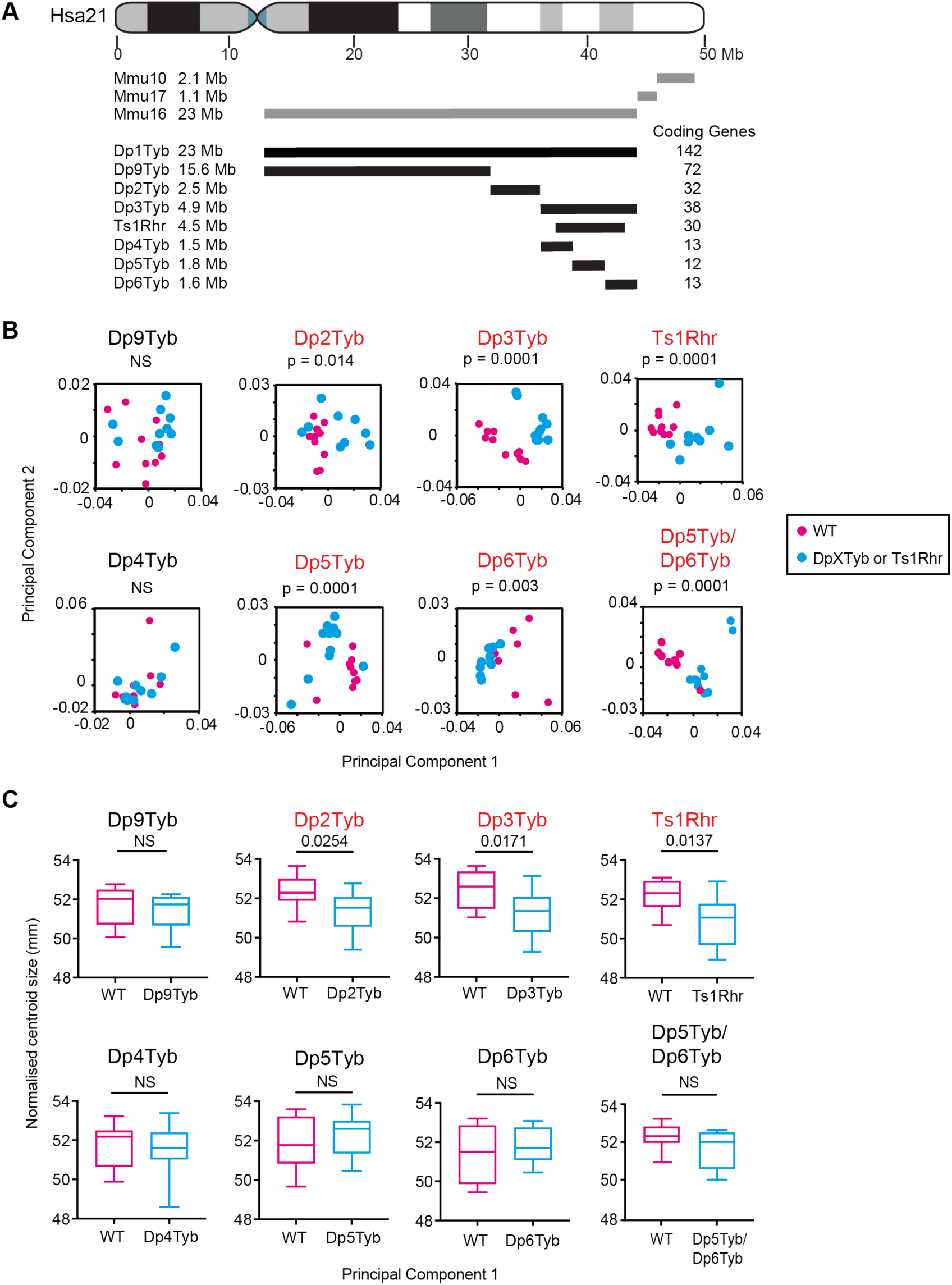
Mapping genes causing cranial dysmorphology in DS. (**A**) Diagram of Hsa21 showing main cytogenic regions (rectangles of different shades) and the centromere (cyan). Regions of orthology to Mmu10, Mmu16 and Mmu17 are indicated in grey lines. Black lines represent duplicated regions in each mouse model, showing the number of coding genes within each region and its size in megabases (Mb). (**B**) PCA (first two components) of Procrustes aligned crania shapes determined using landmark-based morphometrics. Statistical significance (*p*-values) of the difference between WT and mutants was calculated using a multiple permutations test. (**C**) Centroid sizes of crania shown as box and whiskers plots indicating the 25% and 75% centiles (box), range of all data points (whiskers) and the median (line). Statistical significance was calculated using a two-tailed unpaired t-test. N = 10 for each genotype. Mutant strains showing significant differences (*p* < 0.05) are shown in red. NS, not significant (*p* > 0.05).

The region duplicated in Dp1Tyb mice is broken down into three segments in Dp2Tyb, Dp3Tyb and Dp9Tyb mice (Fig. 1A). Of these strains, the crania of Dp2Tyb and Dp3Tyb mice were significantly altered in shape, and decreased in size compared to WT controls, whereas no change was seen in the crania of Dp9Tyb mice (Fig. 1B, C, Table S1). Ts1Rhr mice have a duplication that is entirely contained within the region duplicated in Dp3Tyb mice but is 8 coding genes shorter (Fig. 1A, Fig. S1) (Olson et al., 2004). Ts1Rhr mice showed altered shape and decreased size of the cranium similar to that seen in Dp3Tyb mice (Fig. 1B, C). The Dp4Tyb, Dp5Tyb and Dp6Tyb strains each have a different short duplication that together cover the entire region duplicated in Dp3Tyb mice (Fig. 1A). No abnormality was seen in Dp4Tyb crania while Dp5Tyb and Dp6Tyb crania were altered in shape, although not in size (Fig. 1B, C). Thus, minimally, the cranial changes are caused by one or more genes in each of the Dp2Tyb, Dp5Tyb and Dp6Tyb regions.

Morphometric analysis of the mandibles showed that they were decreased in size in Dp2Tyb and Dp3Tyb mice, but only Dp3Tyb mandibles were altered in shape (Fig. 2A, B, Table S1). Once again, Ts1Rhr mandibles showed changed shape and decreased size, similar to Dp3Tyb mice. The mandibles of Dp9Tyb, Dp4Tyb, Dp5Tyb and Dp6Tyb mice were not significantly affected in either shape or size (Fig. 2A, B). Thus, the genes causing the mandibular phenotype map to the Dp2Tyb and Dp3Tyb regions but disappear once the Dp3Tyb region is broken down further, implying that there must be at least one causative gene in the Dp2Tyb region and at least two in the Dp3Tyb region.

**Figure 2.**
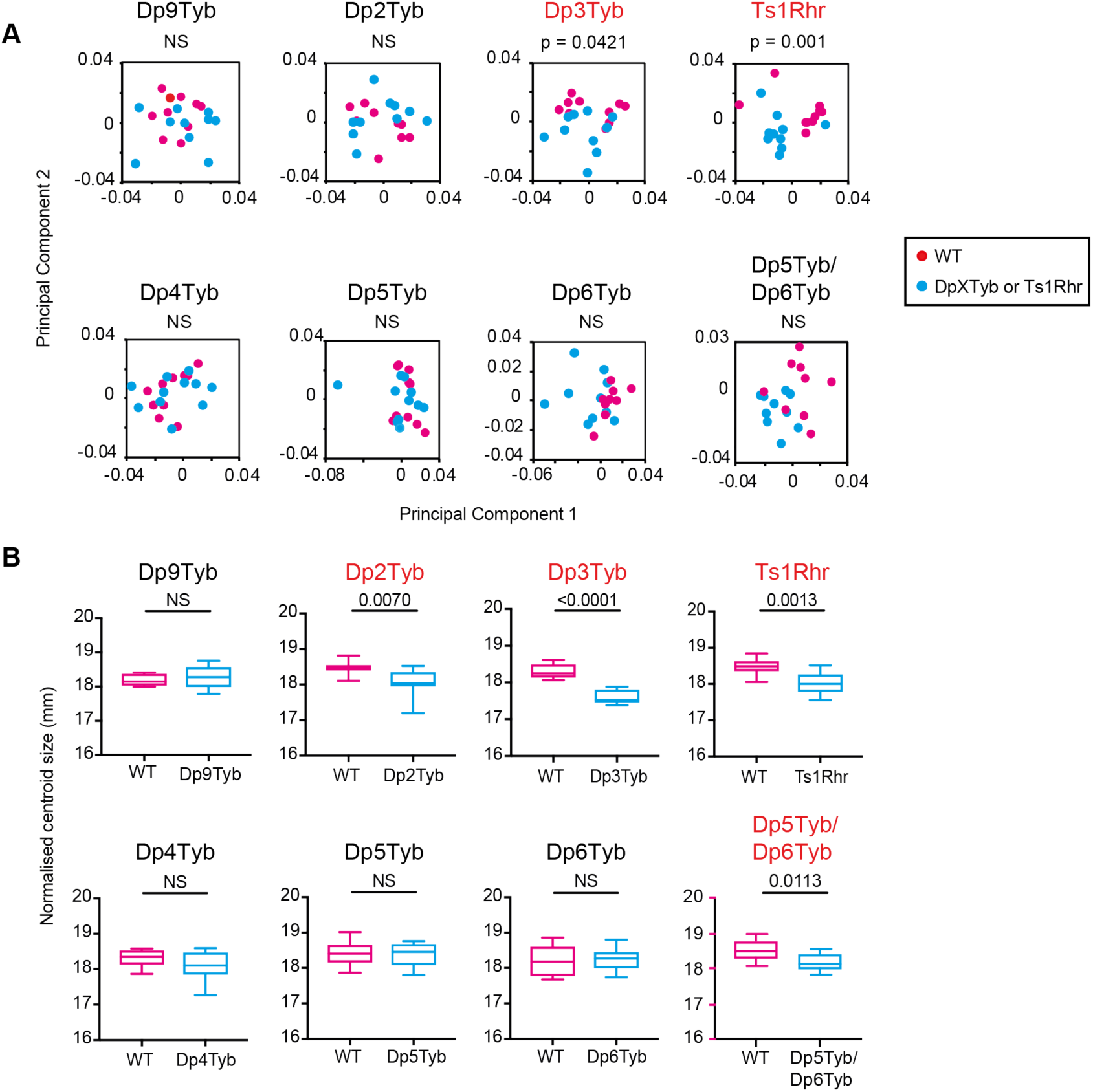
Mapping genes causing mandibular dysmorphology in DS. **(A)** PCA of Procrustes aligned mandible shapes determined using landmark-based morphometrics. Statistical significance was calculated using a multiple permutations test. (**B**) Centroid sizes of mandibles shown as box and whiskers plots indicating the 25% and 75% centiles (box), range of all data points (whiskers) and the median (line). Statistical significance was calculated using a two-tailed unpaired t-test. N = 10 for each genotype. Mutant strains showing significant differences (*p* < 0.05) are shown in red. NS, not significant (*p* > 0.05).

The key aspects of the phenotype of Dp1Tyb crania are midfacial hypoplasia and brachycephaly (Toussaint et al., 2021). To visualise these changes, we compared the location of landmarks on Dp1Tyb crania compared to WT controls (Fig. 3A-C). These showed a posterior shift of anterior landmarks (cyan points in left half of both Fig. 3A and 3B), widening of the skull (see cyan points on the zygomatic arches in the superior view Fig. 3B), a doming of the skull (shown by the superior movement of the cyan points in the lateral view Fig. 3A) and a contraction of the base of the skull (Fig. 3C). Similar changes were seen in Dp2Tyb, Dp3Tyb, Ts1Rhr and Dp5Tyb mice, although they were smaller in magnitude (Fig. 3D-G). In contrast, Dp6Tyb crania showed a ’reverse’ phenotype with a more elongated midface, and a narrower and flatter skull (Fig. 3H, I). Similarly, comparison of the shapes of the mandibles showed that Dp3Tyb and Ts1Rhr had a contraction of the alveolar ramus and condylar process, which was similar to that seen in Dp1Tyb mice, albeit smaller in magnitude (Fig. 3J-L).

**Figure 3.**
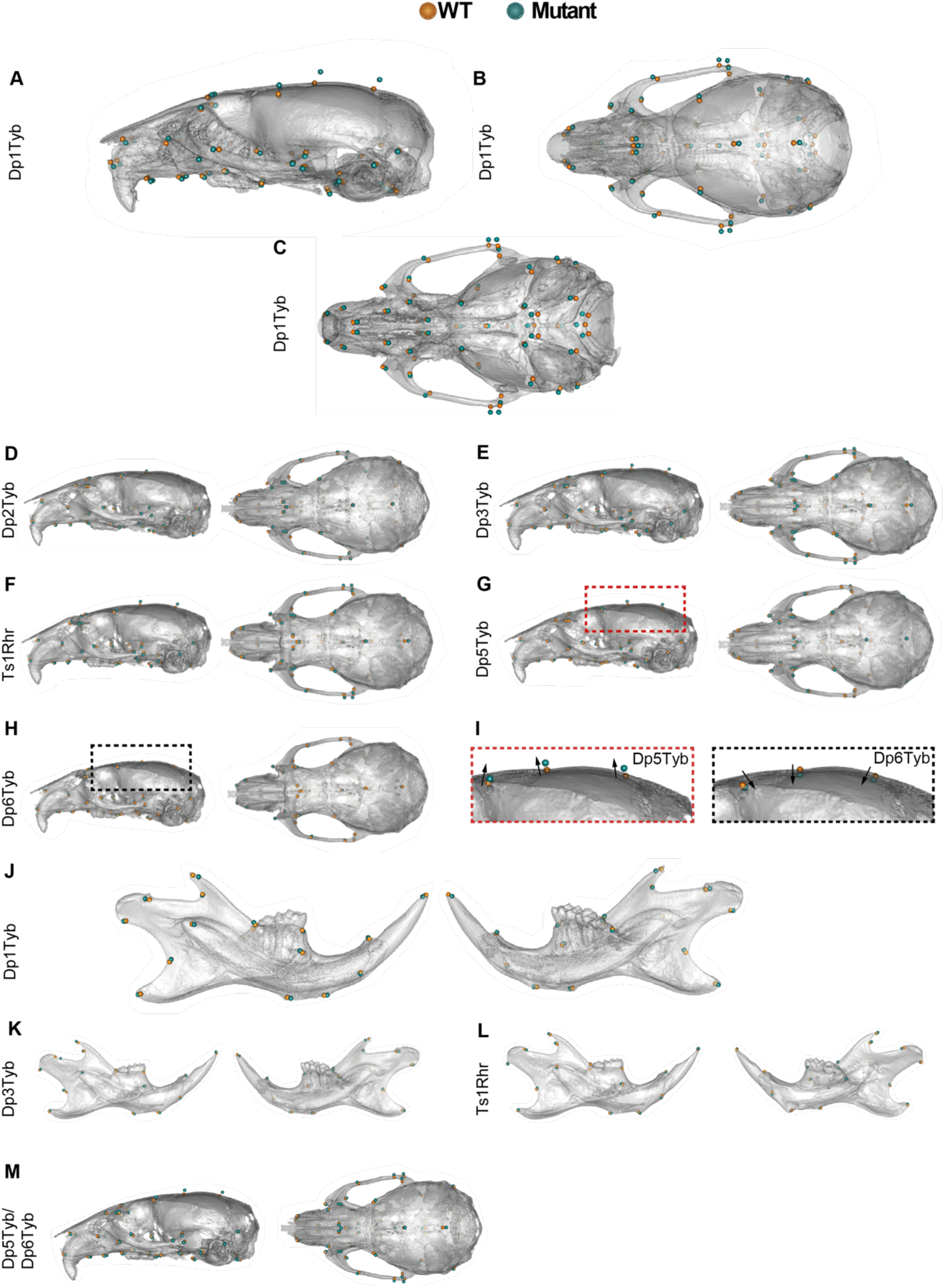
Shape differences in DS mouse model skulls. (**A-C**) Lateral, superior and inferior views of Dp1Tyb skulls adapted from Toussaint et al. (2021). (**D-I**) Mean landmark configurations of WT versus mutant crania with global size differences removed, showing lateral and superior views; zoomed view of cranial vault of Dp5Tyb and Dp6Tyb mice (I), arrows emphasise direction of shape change. (**J-L**) Mean landmark configurations of WT versus mutant mandibles with global size differences removed, showing buccal and lingual views. (**M**) Mean landmark configurations of WT versus Dp5Tyb/Dp6Tyb crania with global size differences removed, showing lateral and superior views.

In the analysis of the three strains that break up the Dp3Tyb region, only Dp5Tyb and Dp6Tyb mice had craniofacial phenotypes, suggesting that the genes causing the Dp3Tyb phenotypes reside in these two regions. To test this, we generated Dp5Tyb/Dp6Tyb double mutant mice and analysed their skulls as above. Compared to WT controls, the crania of Dp5Tyb/Dp6Tyb mice were altered in shape but not size (Fig. 1B, C, Table S1), whereas Dp5Tyb/Dp6Tyb mandibles were smaller but not changed in shape (Fig. 2A, B). The shape phenotype in these mice was similar to the other strains in the mapping panel, with a shortened snout, doming of the skull and relative widening of the zygomatic arches (Fig. 3M). Thus, combining the Dp5Tyb and Dp6Tyb mutations does not fully recapitulate the Dp3Tyb phenotype, implying that here must also be a genetic contribution from the Dp4Tyb region.

To gain an overview of the shape changes across all the mouse strains in the mapping panel, we generated a PCA plot by directly comparing the landmark data for the crania from all mouse lines. The plot was normalised by placing the means of all the WT cohorts at the origin, allowing the direction and magnitude of the shape change of each mutant cohort to be directly compared (Fig. 4A). The DS-like cranial shape changes primarily separate the strains along principal component 1, with Dp1Tyb mice showing the largest change. Dp3Tyb and Ts1Rhr crania show a shape change in the same direction as Dp1Tyb, but smaller in magnitude. Leave-one-out cross-validation analysis shows that PCA is unable to robustly distinguish the two strains, implying that the additional 8 genes that are in three copies in Dp3Tyb mice but not Ts1Rhr mice (Fig. S1) are not contributing significantly to phenotype. Dp5Tyb mice show a small shape change in the same direction as Dp3Tyb and Dp1Tyb mice, which is increased in the Dp5Tyb/Dp6Tyb double mutant but is still not as large as the change in Dp3Tyb mice, implying that there must also be causative genes in the Dp4Tyb region. Dp2Tyb mice also show a small phenotypic change in the direction of Dp1Tyb mice, suggesting that this region may contain one or more causative genes that together with the causative genes in the Dp3Tyb region give rise to the full Dp1Tyb phenotype. Finally, Dp6Tyb mice show a reverse phenotype in principal component 1, visible as a shape change in the opposite direction in many respects to the other strains. A summary of the phenotype mapping seen across all the strains is shown in Fig. 4B. Taken together, these results imply that the Dp1Tyb dysmorphology must be caused by increased dosage of at least four genes, one in each of the regions duplicated in Dp2Tyb, Dp4Tyb, Dp5Tyb and Dp6Tyb mice (Fig. S1). Furthermore, the causative genes in the Dp4Tyb and Dp6Tyb regions most likely lie within the overlap with the region duplicated in Ts1Rhr, excluding the 8 genes that are duplicated in Dp3Tyb but not Ts1Rhr.

**Figure 4.**
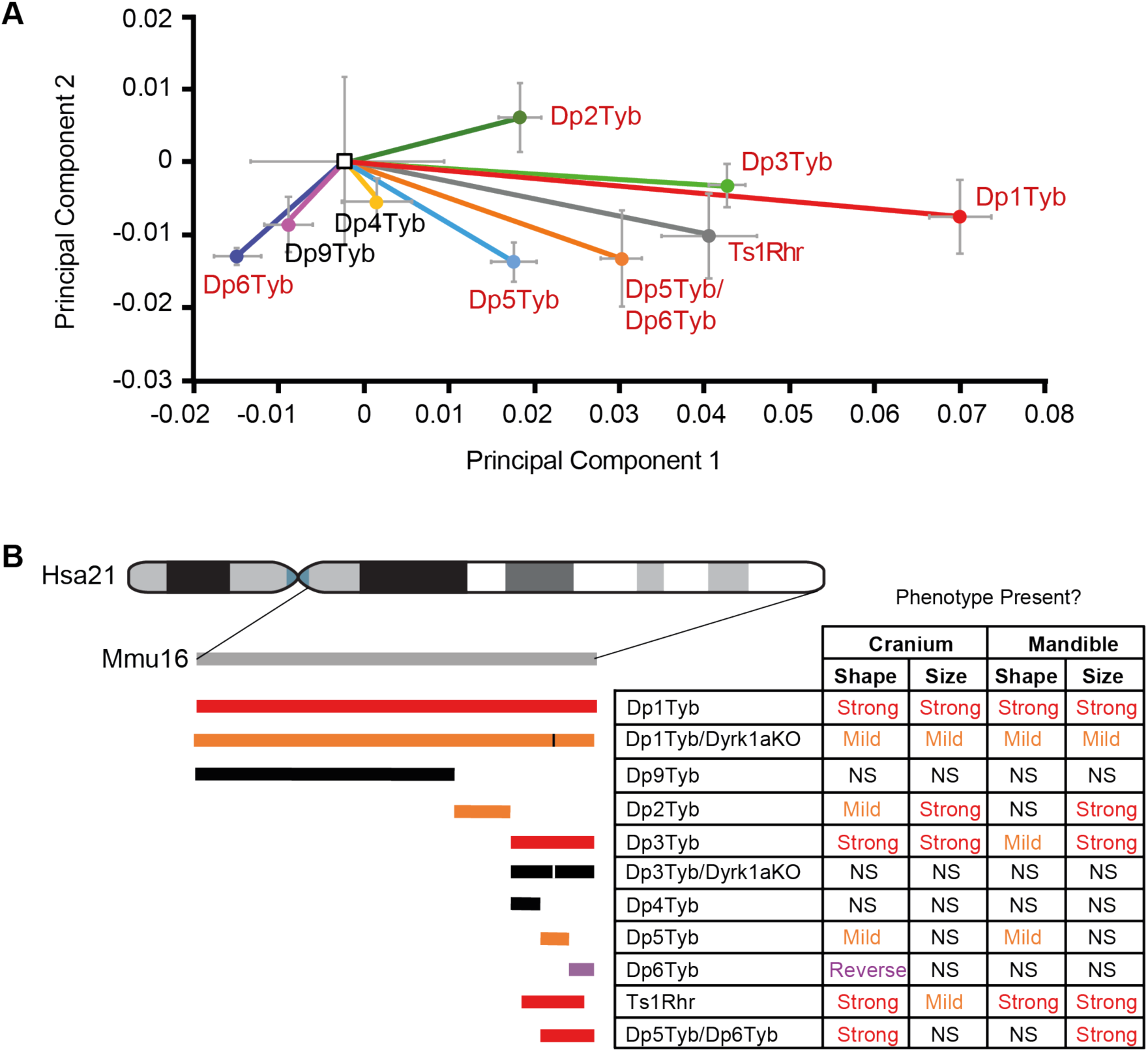
Increased dosage of at least four genetic loci contribute to DS craniofacial dysmorphology. (**A**) Normalised PCA plot of cranial shapes of the indicated mutant mouse strains. Plots have been normalised to place the mean PCA values for each WT cohort at the origin, with vectors indicating the mean PCA values (± standard deviation) of each mutant strain. Mutant strains showing significant differences (*p* < 0.05) compared to their WT controls are shown in red. (**B**) Summary of craniofacial phenotypes in the genetic mapping panel. Diagram of duplicated regions on the left as in Figure 1A, with the addition of the Dp5Tyb/Dp6Tyb, Dp1Tyb/Dyrk1aKO and Dp3Tyb/Dyrk1aKO strains. Vertical lines indicate the location of the disrupted *Dyrk1a* gene. Duplicated regions are coloured to indicate severity of the cranial shape phenotype: red, severe; orange mild; purple, reversed phenotype in Dp6Tyb; black, no phenotype. Table shows whether cranial and mandibular phenotypes are present in the strains. NS, not significant (*p* > 0.05).

### *Dyrk1a* is one of the genes required in three copies to cause craniofacial dysmorphology in Dp1Tyb mice

The strain with the smallest duplicated region to show a craniofacial phenotype is Dp5Tyb, which contains an extra copy of just 12 coding genes. One of these genes, *Dyrk1a*, which codes for the DYRK1A serine/threonine protein kinase, has been shown to contribute to several DS phenotypes when present in three copies (Ahn et al., 2006; Altafaj et al., 2001; Brault et al., 2021; Duchon et al., 2021; Jiang et al., 2015; London et al., 2018; McElyea et al., 2016; Souchet et al., 2014; Thomazeau et al., 2014; Watson-Scales et al., 2018; Yin et al., 2017). Thus, we tested the requirement for three copies of *Dyrk1a* for the craniofacial phenotypes of Dp1Tyb and Dp3Tyb mice. We crossed mice with a loss of function *Dyrk1a* allele (*Dyrk1a*^+/-^) to Dp1Tyb and Dp3Tyb mice to generate Dp1Tyb/Dyrk1aKO and Dp3Tyb/Dyrk1aKO mice in which one of the three copies of *Dyrk1a* was inactivated leaving 2 functional copies, while maintaining all other duplicated genes at three copies. Morphometric analysis of Dp1Tyb/Dyrk1aKO skulls showed that their crania and mandibles were altered in shape compared to WT controls, but were also significantly different to Dp1Tyb crania, lying between WT and Dp1Tyb mice on a PCA plot (Fig. 5A, B). Similarly, Dp1Tyb/Dyrk1aKO crania and mandibles were reduced in size, but the change was not as large as that seen in Dp1Tyb mice (Table S1). Analysis of Dp3Tyb/Dyrk1aKO crania and mandibles showed that they were not significantly different from WT controls in either shape or size (Fig. 5C, D, Table S1). Thus, reducing *Dyrk1a* copy number from three to two partially rescues the Dp1Tyb craniofacial dysmorphology, and fully rescues the Dp3Tyb phenotype, demonstrating that *Dyrk1a* is one of the causative genes (Fig. 4B, S1).

**Figure 5.**
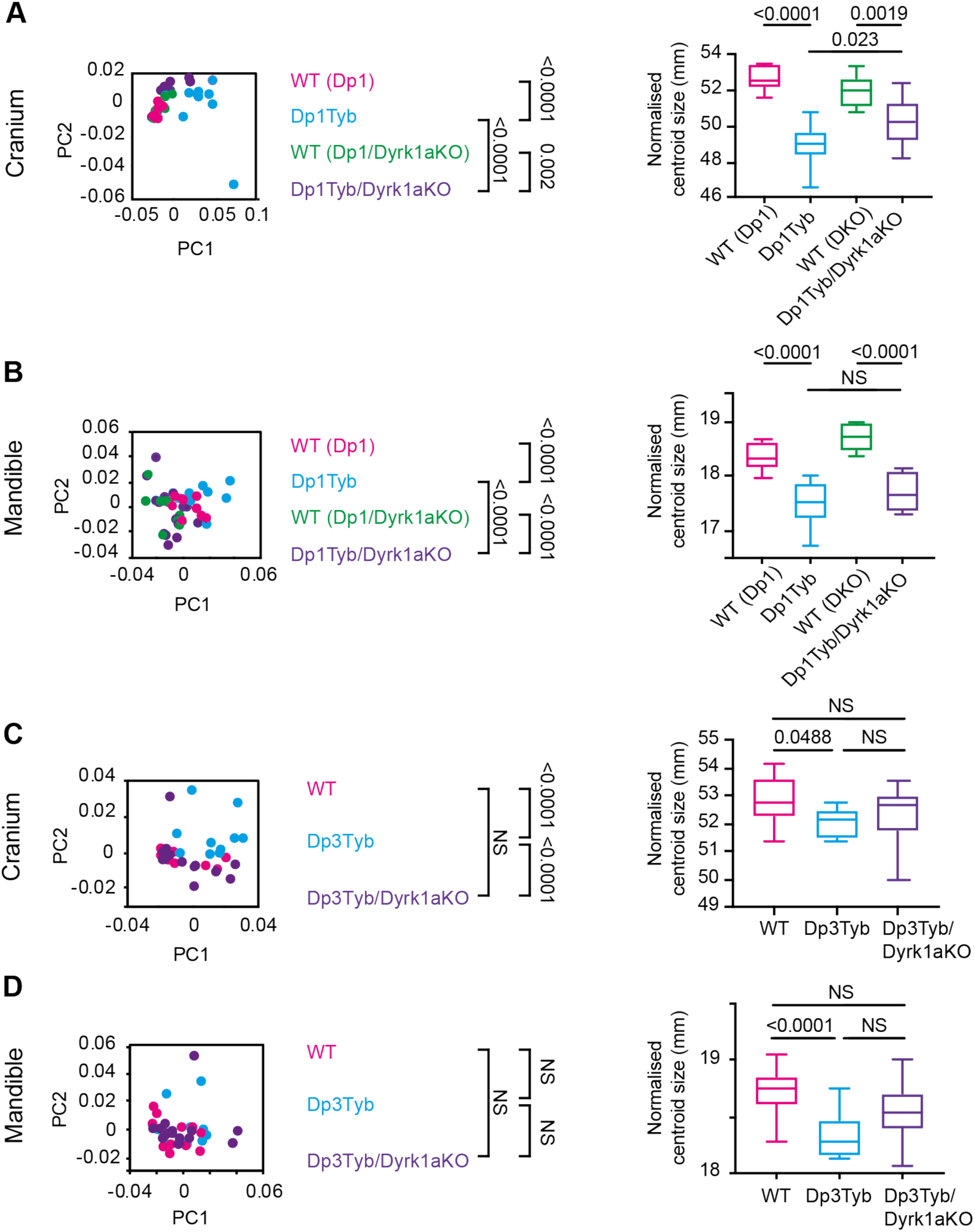
Analysis of Dp5Tyb/Dp6Tyb and Dp1Tyb/Dyrk1aKO mice. (**A-D**) PCA of Procrustes aligned shapes determined using landmark-based morphometrics and centroid sizes of crania (A, C) and mandibles (B, D) of Dp1Tyb and Dp1Tyb/Dyrk1aKO mice, each with its own WT control mice as indicated in parentheses (A, B) and Dp3Tyb, Dp3Tyb/Dyrk1aKO and WT control mice (C, D). Centroid size shown as box and whiskers plots indicating the 25% and 75% centiles (box), range of all data points (whiskers) and the median (line). Statistical significance (*p*-values) was calculated using a multiple permutations test for PCA plots and a two-tailed unpaired t-test for centroid sizes. N = 10 for each genotype. Dp1Tyb data in A and B from Toussaint et al. (2021). NS, not significant (*p* > 0.05).

### Aberrant synchondroses in Dp1Tyb skulls

When studying the 16 week-old Dp1Tyb mice we observed that cartilaginous growth points in the base of the cranium known as synchondroses were often dysmorphic. In mice, there are two midline synchondroses: the more anterior, intersphenoid synchondrosis (ISS) (Fig. 6A, green arrow) and the more posterior, spheno-occipital synchondrosis (SOS) (Fig. 6A, purple arrow). Using both volumetric reconstructions and digital slices of the base of the skull, we classified ISS morphology in each animal as normal, partially dysmorphic or fully fused where no cartilaginous gap remained (Fig. 6A). Similarly, SOS morphology was classified as normal or partially dysmorphic. Dp1Tyb mice had a significantly increased frequency (> 50%) of dysmorphic ISS and SOS (Fig. 6B, C). Analysis of synchondroses across the mapping panel showed that only Dp3Tyb and Ts1Rhr mice had significantly increased rates of dysmorphic ISS, although several other strains (Dp2Tyb, Dp5Tyb and Dp5Tyb/Dp6Tyb) showed evidence of dysmorphology which did not reach significance (Fig. 6B). Only Dp1Tyb mice had aberrant SOS (Fig 6C). Disruption of one copy of *Dyrk1a* caused a reduction in the severity of ISS dysmorphology in Dp1Tyb mice and eliminated it in Dp3Tyb mice, and resulted in rescue of the SOS dysmorphology in Dp1Tyb mice (Fig. 6B, C). Thus, three copies of *Dyrk1a* are required for dysmorphic synchondroses. Taken together, the frequency of dysmorphic synchondroses tracked closely with cranial shape changes across the mapping panel, suggesting they may be caused by related pathological mechanisms (Fig. 6D).

**Figure 6.**
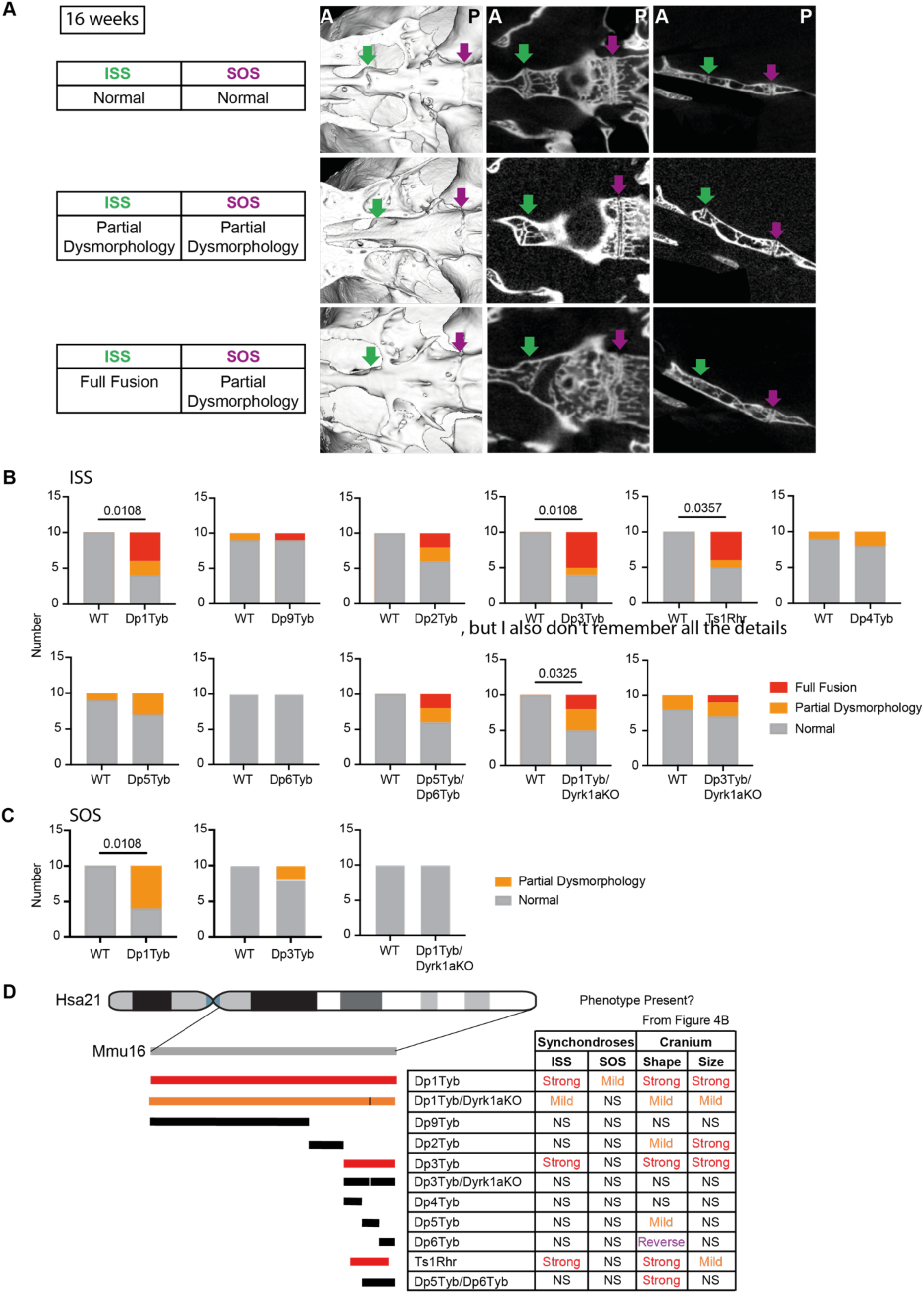
DS mouse models have aberrant synchondroses. (**A**) Inferior volumetric rendering (left), horizontal 2D slice (middle) and sagittal slice (right) of crania at 16 weeks of age, showing representative examples of synchondroses with normal, partially dysmorphic and fully fused phenotypes; green arrows, ISS; magenta arrows, SOS. Note dysmorphic SOS are shown in both the partial ISS dysmorphology and full ISS fusion images. A, anterior; P, posterior. (**B, C**) Incidence of fully fused, partially dysmorphic and normal ISS (B) and SOS (C) in indicated mutant mouse strains. Only Dp1Tyb and Dp3Tyb are shown for SOS morphology because all other strains in the mapping panel showed no SOS dysmorphology. Statistical significance was calculated using Fisher’s exact test, indicating significant differences (*p* < 0.05). Where no *p*-value is shown, comparison of mutants to WT is not significant (*p* > 0.05). N = 10 for each genotype. (**D**) Summary of synchondroses phenotypes in the DS genetic mapping panel as in Figure 4B. Colours of duplicated regions indicate severity of the ISS phenotype. Table shows whether ISS and SOS phenotypes are present in the strains, with the table of cranial phenotypes repeated from Figure 4B to allow easier comparison. NS, not significant (*p* > 0.05).

### Developmental trajectory of craniofacial dysmorphology in Dp1Tyb mice

To gain a better mechanistic understanding of the origins of the craniofacial defects in Dp1Tyb mice, we analysed the skulls of young mice at 21 days post-partum (P21) by µCT and landmark-free morphometrics (Toussaint et al., 2021). We chose to use a landmark-free method for the analysis, because the changes at this age were more subtle than at 16 weeks, and thus more readily captured with landmark-free morphometrics compared to a landmark-based approach. The shape of the Dp1Tyb cranium was significantly different to WT controls, but the mandible shape was not affected and there were no significant differences in size of either cranium or mandible (Fig. 7A, B, Video S1). Qualitatively, the cranial shape difference at this stage was similar to that observed at 16 weeks (Fig. 3A) (Toussaint et al., 2021). The snout was displaced towards the posterior and the calvaria were displaced laterally as seen in the displacement heat maps (Fig. 7C). This is further confirmed by the stretch heatmaps (Toussaint et al., 2021) which indicate that the snout is contracted and the calvaria are expanded as seen in the blue and red regions of Fig. 7D, respectively. We also examined the mid-line synchondroses in the cranial base. In WT control mice these were still patent, with no mineralisation (Fig. 7E, F), as expected based on previous analysis of C57BL/6J mice at P21 (Vora et al., 2016). In contrast, many Dp1Tyb mice had partially or fully fused ISS, with evidence of aberrant mineralisation bridges (Fig. 7E, F).

**Figure 7.**
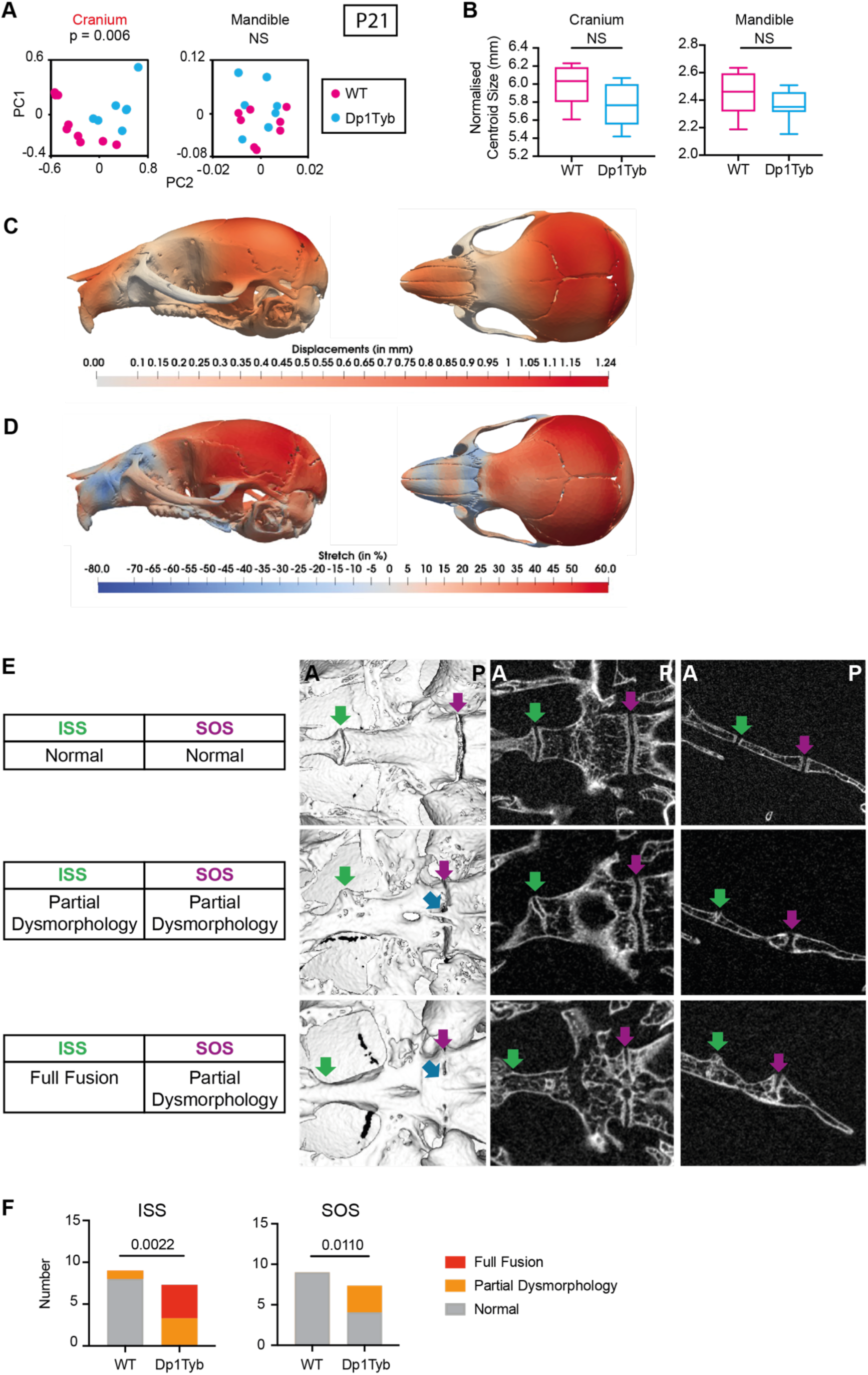
Craniofacial dysmorphology in Dp1Tyb mice at P21. (**A, B**) PCA of Procrustes aligned crania and mandible shapes determined using landmark-free morphometrics (A) and centroid sizes of crania and mandibles (B) of P21 Dp1Tyb and WT control mice. Centroid size data shown as box and whiskers plots indicating the 25% and 75% centiles (box), range of all data points (whiskers) and the median (line). (**C, D**) Heatmaps of displacement (C) and stretch (D) of Dp1Tyb P21 crania compared to WT controls, showing lateral and superior views. See also Video S1. (**E**) Inferior and lateral views of volumetric reconstructions of crania at P21, showing representative examples of synchondroses with normal, partially dysmorphic and fully fused phenotypes; green arrows, ISS; magenta arrows, SOS; blue arrows, mineralisation bridges. Note dysmorphic SOS are shown in both the partial ISS dysmorphology and full ISS fusion images. A, anterior; P, posterior. (**F**) Incidence of fully fused, partially dysmorphic and normal ISS and SOS. Statistical significance (*p*-values) was calculated using a multiple permutations test (A), a two-tailed unpaired t-test (B) and Fisher’s exact test (F). WT N = 9; Dp1Tyb N = 7. NS, not significant (*p* > 0.05).

To extend this analysis further back in development, we studied the skulls of Dp1Tyb embryos at embryonic day 16.5 (E16.5) and E18.5 using µCT and landmark-free morphometrics (Toussaint et al., 2021), which allowed us to compare the developing cranial bones, which lack many of the usual landmarks. Crucially, it also allows analysis of local expansion or contraction (“stretch”) rather than net displacement. We found that the crania of Dp1Tyb embryos at both E16.5 and E18.5 were significantly altered in shape (Fig. 8A, C, Video S2), but there was no significant change in overall size (Fig. 8B, D). Examination of the shape changes showed that at E18.5 Dp1Tyb crania are flatter (Fig. 8E, F, + marks, Video S2) which is opposite to the domed phenotype seen at P21 and 16 weeks. However, like the postnatal stages, E18.5 Dp1Tyb crania had a contraction of the snout (Fig. 8F, blue region on the left-hand side). At E16.5 Dp1Tyb crania showed a reduction in the size of the bones in the front half of the cranium (Fig. 8G, red regions, Fig. 8H, blue regions) and the zygomatic arch (Fig. 8G, H, asterisks). Strikingly, the E16.5 Dp1Tyb embryos also showed a relative expansion of the basioccipital and occipital condyle bones (Fig. 8H, black arrow and red region). Similar analysis of Dp1Tyb/Dyrk1aKO embryos showed that their crania were not significantly different from WT in shape or size at either E16.5 or E18.5 (Fig. 8A-D). Thus, the changes in cranial shape of Dp1Tyb embryos are already visible at E16.5 and are dependent on three copies of *Dyrk1a*.

**Figure 8.**
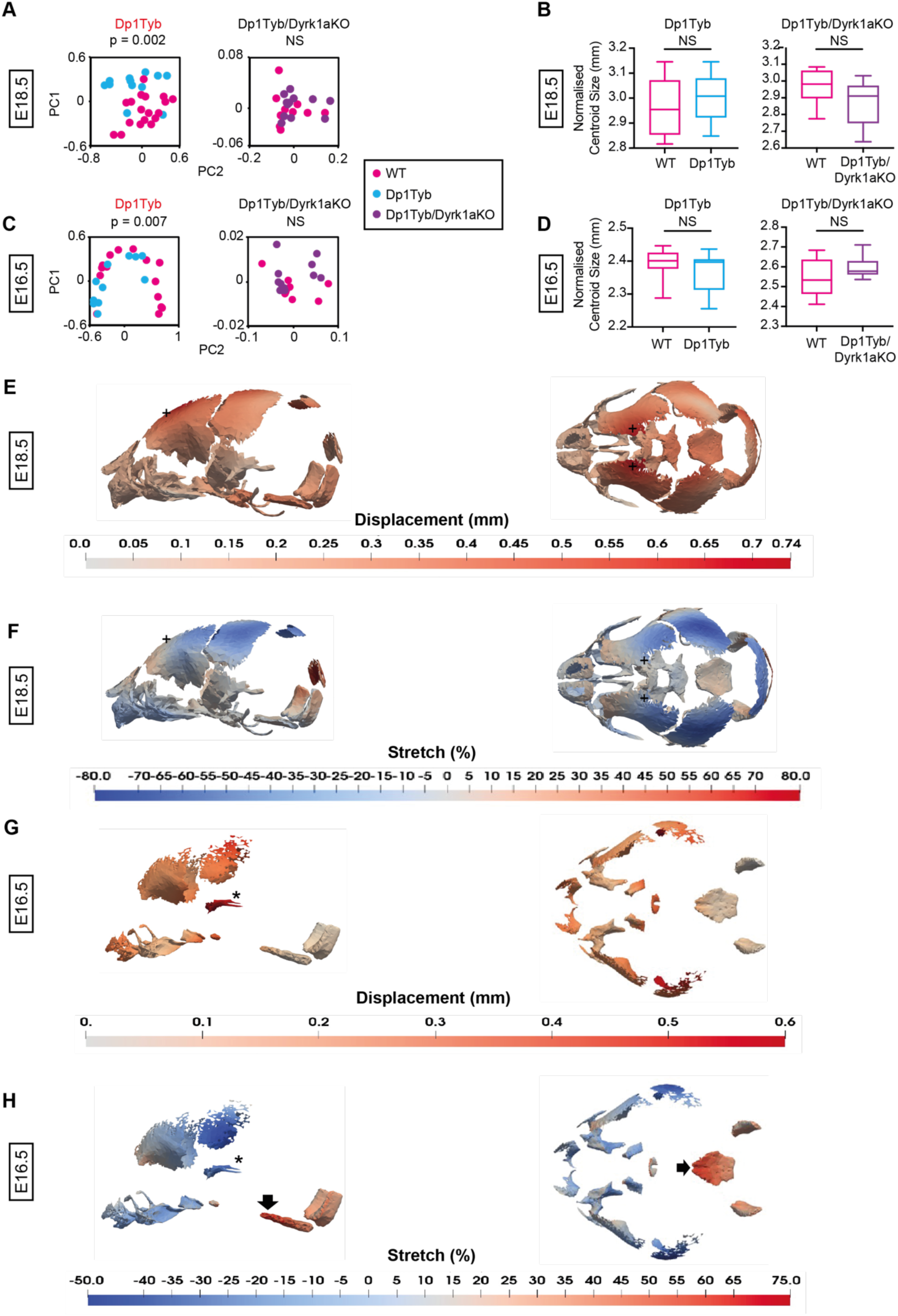
*Dyrk1a*-dependent craniofacial dysmorphology in Dp1Tyb embryos at E18.5 and E16.5. (**A-D**) PCA of Procrustes aligned crania shapes determined using landmark-free morphometrics (A, C) and centroid sizes (B, D) of Dp1Tyb and Dp1Tyb/Dyrk1aKO embryos and their respective WT controls at E18.5 (A, B) and E16.5 (C, D). Centroid size data shown as box and whiskers plots indicating the 25% and 75% centiles (box), range of all data points (whiskers) and the median (line). Statistical significance (*p*-values) was calculated using a multiple permutations test for PCA plots and a two-tailed unpaired t-test for centroid sizes. At E18.5 WT (Dp1Tyb) N = 18; Dp1Tyb N = 12; WT (Dp1Tyb/Dyrk1aKO) N = 10, Dp1Tyb/Dyrk1aKO N = 12. At E16.5 WT (Dp1Tyb) N = 15; Dp1Tyb N = 11; WT (Dp1Tyb/Dyrk1aKO) N = 10; Dp1Tyb/Dyrk1aKO N = 11. (**E-H**) Heatmaps of displacement (E, G) and stretch (F, H) of Dp1Tyb crania compared to WT controls at E18.5 (E, F) and E16.5 (G, H), showing lateral and superior views. See also Video S2. Key: +, frontal bone (panels E, F); **\***, zygomatic arch (panels G, H); black arrow, basioccipital bone (panel H). NS, not significant (*p* > 0.05).

### Increased dosage of *Dyrk1a* causes decreased size and reduced differentiation of frontal bone primordia

Morphometric analysis of Dp1Tyb mice at several stages indicated that the most severely affected bones of the skull were those of neural crest (NC) origin i.e., the frontal and facial bones and the ISS (Jiang et al., 2002; McBratney-Owen et al., 2008) with relatively modest abnormality in skull bones of mesodermal origin (the occipital and basioccipital) (Fig. 8H). The apparent size reduction in the NC-derived bones could be due to fewer cells being present or less µCT-detectable mineralisation, since the latter expands from the centres of the bone primordia towards their edges. To determine which of these was the case, we focussed on analysing development of one of these NC-derived structures, the frontal bones. Since a phenotype was already present in the frontal bones of Dp1Tyb embryos at early stages of ossification (E16.5), we assessed whether changes could be detected earlier in development by analysing the frontal bone primordia, a mesenchymal condensation of NC origin, at E13.5 (Fig. 9A). The cells within these primordia will undergo differentiation into osteoblasts that produce mineralised bone. Histological analysis showed that frontal bone primordia of Dp1Tyb E13.5 embryos were reduced in size and had fewer cells (Fig. 9A, B). Notably, this defect was not seen in Dp1Tyb/Dyrk1aKO embryos (Fig. 9A, B). Thus, three copies of *Dyrk1a* are required for the decreased size and cellularity of frontal bone primordia in Dp1Tyb embryos.

**Figure 9.**
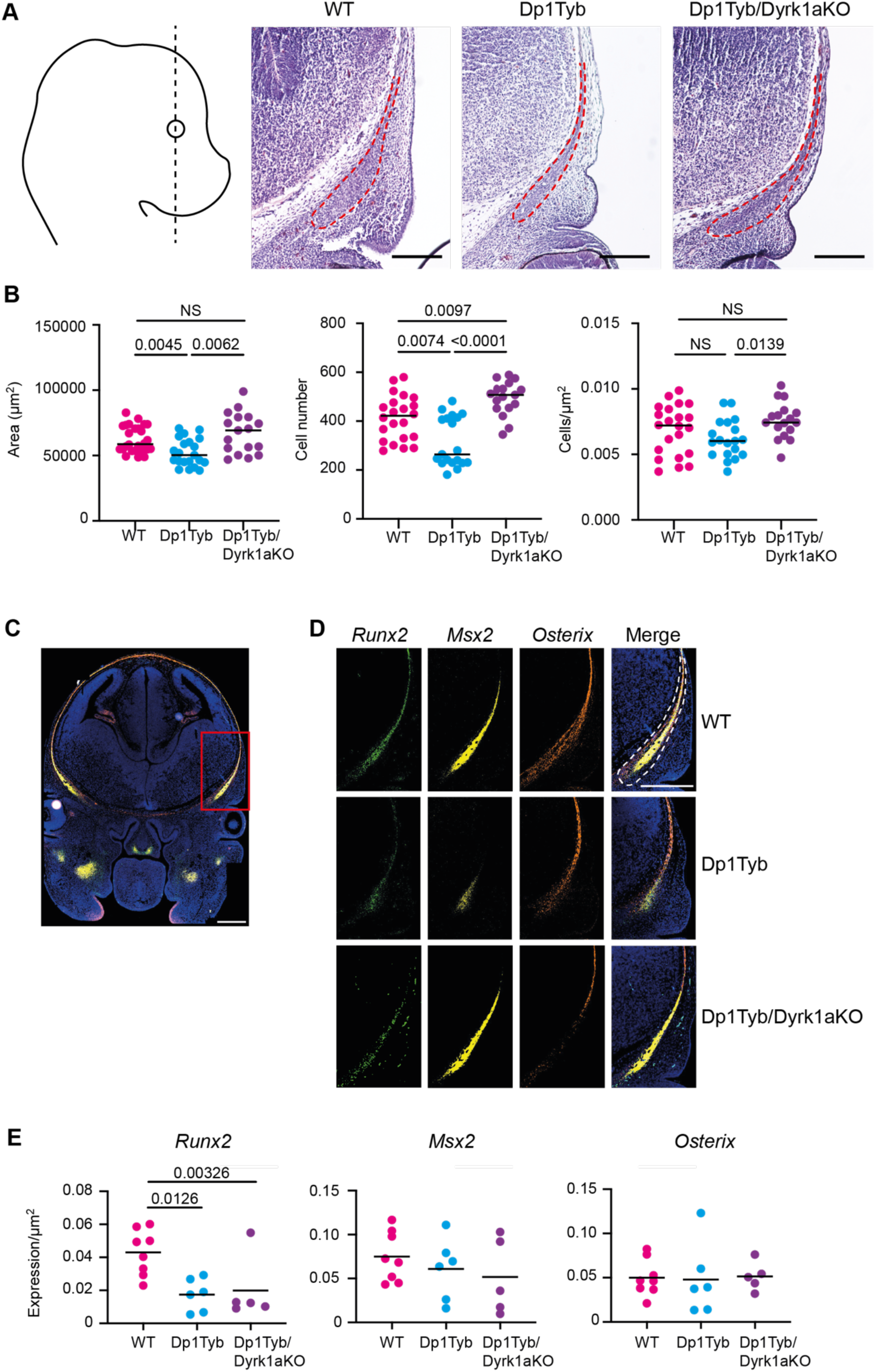
Increased dosage of *Dyrk1a* causes decreased size of frontal bone primordia. (**A**) Representative H&E images of frontal bone primordia (dashed red line) of E13.5 embryos of the indicated genotypes sectioned coronally as shown by the dashed black line on diagram showing a lateral view of the embryo. (**B**) Area, cell number and cell density of coronal frontal primordia. Each dot is a different embryo, black line indicates mean. (**C, D**) Representative RNAscope images of coronal sections through the heads of E13.5 embryos in the same plane as in A, showing hybridisation signals for *Runx2* (green), *Msx2* (yellow) and *Osterix* (orange) mRNA expression either merged (C), or each gene separately and merged (D). The section in C is through a WT embryo showing the whole head and the red box indicates the area of high magnification shown in D. Sections in D are from embryos of the indicated genotypes. Dashed white line indicates the frontal bone primordium. Sections were stained with DAPI to visualise nuclei. (**E**) Mean of *Runx2*, *Msx2* and *Osterix* mRNA expression in frontal bone primordia of E13.5 embryos of the indicated genotypes. Each dot is a different embryo, black line indicates mean. Statistical significance calculated with Tukey’s multiple comparison test. All scale bars are 200 µm.

Another mechanism which could lead to aberrant facial bones in Dp1Tyb embryos is altered ossification. To investigate this, we used RNAscope, a type of *in situ* hybridisation, on sections of the frontal bone primordia at E13.5 to quantify the expression of *Runx2*, *Msx2* and *Osterix*, genes that are critical for the differentiation of mesenchymal cells into the osteogenic lineage (Han et al., 2007; Maeno et al., 2011; Matsubara et al., 2008; McGee-Lawrence et al., 2014; Nakashima et al., 2002) (Fig. 9C, D). The expression of the genes was normalised to area, to account for the reduced size of the primordia in Dp1Tyb embryos. We found that the expression of *Runx2*, but not *Msx2* or *Osterix*, was reduced in Dp1Tyb primordia, suggesting that there may be ossification defects in the mutant embryos (Fig. 9E). Interestingly, reduced *Runx2* expression was still present in Dp1Tyb/Dyrk1aKO embryos, thus this phenotype is caused by three copies of genes other than *Dyrk1a* (Fig. 9E).

### Increased dosage of *Dyrk1a* causes reduced proliferation of cranial neural crest cells

The decrease in the size and cell number of Dp1Tyb frontal bone primordia could be due to defects in different processes. NC cells, the progenitors of this structure, undergo a key migratory phase early in their development and are also highly proliferative (Wu et al., 2017). A defect in either of these cellular behaviours could lead to the observed decrease in area and cellularity of the Dp1Tyb primordia, thus we examined both processes. To measure migration, explants of E8.5 anterior neural tubes were cultured and individual migrating cells tracked for 18 h (Fig. 10A). We found that Dp1Tyb NC cells had no significant change in speed or persistence (straightness) of migration compared to WT cells (Fig. 10B), suggesting that early defects in NC migration are unlikely to account for the smaller frontal bone primordia in the mutant embryos.

**Figure 10.**
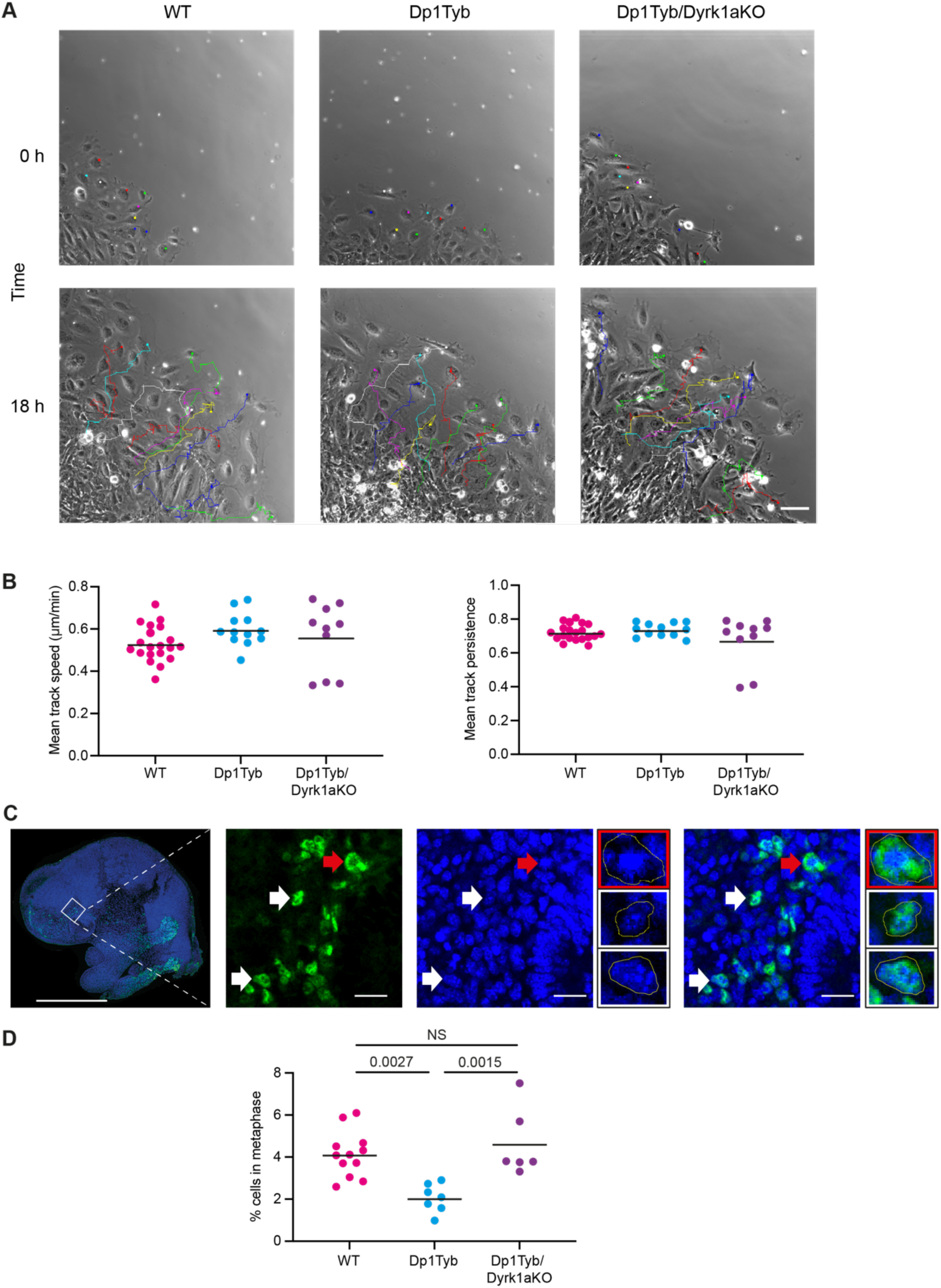
Increased dosage of *Dyrk1a* causes reduced proliferation of cranial neural crest cells. (**A**) Representative first and last frames of 18-hour time lapse videos of tracked neural crest cells migrating out from a E8.5 neural tube explant of the indicated genotypes. Coloured dots in the first frame indicate cells whose migration is tracked and shown in the last frame. Scale bar, 100 µm (**B**) Mean track speed and mean track persistence (straightness) of migrating neural crest cells from E8.5 neural tube explants of the indicated genotypes. Each dot is a different embryo, black line indicates mean. (**C**) Representative low magnification image (left) and higher magnification views (other three large images) of a parasagittal section through a E9.5 mouse embryo head stained for SOX10 (green) and DAPI (blue) in the facial prominence. White and red arrows indicate examples of SOX10^+^ cells that are not in metaphase or in metaphase, respectively, and these same cells are shown enlarged in the three small images, showing the SOX10^+^ cell in metaphase (red border) and two cells not in metaphase (white border). Yellow lines indicate cell outlines. Scale bar is 1 mm (left-hand image) and 200 µm (other three large images). (**D**) Percentage of SOX10^+^ cells that are in metaphase in the facial prominence of E9.5 embryos calculated from images such as those in C. Each dot is a different embryo, black line indicates mean. Statistical significance calculated with Tukey’s multiple comparison test.

Finally, we examined the proliferation of NC-derived cells in the facial prominence of E9.5 embryos by imaging these structures for SOX10 expression to identify NC-derived cells and DAPI to visualise cells in metaphase (Fig. 10C). We found that Dp1Tyb embryos had many fewer SOX10+ cells in metaphase compared to WT controls, implying that the Dp1Tyb mutation causes reduced proliferation of these cells, which may account for the reduced size and cellularity of the frontal bone primordia (Fig. 10D). Importantly, there was no proliferation deficit in Dp1Tyb/Dyrk1aKO embryos, demonstrating that increased dosage of *Dyrk1a* was required for reduced proliferation of NC-derived cells in the facial prominence.

Taken together our results show that craniofacial dysmorphology in Dp1Tyb mice is caused by increased dosage of at least four genes, one of which is *Dyrk1a*. Furthermore, we demonstrate that a third copy of *Dyrk1a* results in decreased proliferation of the NC-derived cells that give rise to frontal bone primordia. This may account for the altered sizes of the frontal bones and, ultimately, skull dysmorphology.

## DISCUSSION

DS is the most common cause of craniofacial dysmorphology in humans, yet the genetic and developmental mechanisms that underly it are poorly understood. Taking advantage of the Dp1Tyb mouse model of DS with three copies of a 142-gene Hsa21-orthologous region of Mmu16 (Lana-Elola et al., 2016) and a DS-like craniofacial phenotype (Toussaint et al., 2021), we used genetic mapping to identify 4 shorter regions (A to D) of Mmu16 that each contain one or more dosage-sensitive genes required in three copies to cause the craniofacial dysmorphology (Fig. S1). Region A contains 32 coding genes duplicated in Dp2Tyb mice, region B contains 11 genes duplicated in Dp4Tyb but not Ts1Rhr mice, region C contains 12 genes duplicated in Dp5Tyb mice and region D contains 7 genes duplicated in Dp6Tyb but not Ts1Rhr mice.

We demonstrated that *Dyrk1a* is one of the causative genes in region C, since reducing the copy number of *Dyrk1a* from three to two partially rescues the craniofacial phenotype of Dp1Tyb mice and fully rescues the phenotype of Dp3Tyb mice. DYRK1A is a protein kinase implicated in multiple DS phenotypes, which may be acting through several mechanisms (Ahn et al., 2006; Altafaj et al., 2001; Brault et al., 2021; Duchon et al., 2021; Jiang et al., 2015; London et al., 2018; McElyea et al., 2016; Souchet et al., 2014; Thomazeau et al., 2014; Watson-Scales et al., 2018; Yin et al., 2017). DYRK1A inhibits the function of NFAT transcription factors by phosphorylating them and thereby promoting their exit from the nucleus (Arron et al., 2006). Since NFAT proteins regulate the differentiation of both osteoblasts and osteoclasts, (Aliprantis et al., 2008; Koga et al., 2005; Lee et al., 2009; Winslow et al., 2006) and mice deficient in NFATC2 and NFATC4 have brachycephaly and mid-facial hypoplasia (Arron et al., 2006), increased dosage of *Dyrk1a* in Dp1Tyb mice may be promoting the craniofacial defects by inhibiting NFAT function and hence perturbing osteogenesis.

Alternatively, overexpression of *Dyrk1a* decreases cell proliferation (Liu et al., 2014; Yabut et al., 2010), while inhibition of DYRK1A leads to increased proliferation (Dirice et al., 2016; Shen et al., 2015). DYRK1A may affect proliferation by regulating Cyclin D1, a key regulator of the cell cycle (Sherr, 1995; Yang et al., 2006). DYRK1A phosphorylates Cyclin D1 on threonine 286 causing its degradation by the proteasome, thereby impairing progression from G1 to S phases of the cell cycle (Chen et al., 2013; Soppa et al., 2014). Thus, increased dosage of *Dyrk1a* in Dp1Tyb mice may lead to decreased proliferation which could contribute to craniofacial defects. In support of this, we measured decreased proliferation of NC-derived cells in the frontal prominence of Dp1Tyb embryos, a phenotype which was dependent on three copies of *Dyrk1a*.

Other plausible causative genes include *Rcan1* in region A, *Morc3*, *Sim2* and *Ttc3* in region B and *Dscam* in region D. RCAN1 is an inhibitor of calcineurin, a phosphatase which dephosphorylates NFAT proteins promoting their nuclear entry. Thus, increased RCAN1 expression may cooperate with increased DYRK1A to inhibit NFAT function and thus impair osteogenesis (Arron et al., 2006). MORC3 is a nuclear protein which regulates osteoclast and osteoblast formation (Jadhav et al., 2016), hence increased dosage of MORC3 in Dp1Tyb mice could also perturb osteogenesis. Mice deficient in the SIM2 transcription factor have craniofacial abnormalities, indicating that SIM2 plays an important role in craniofacial development (Shamblott et al., 2002). TTC3 targets AKT for ubiquitination and reduces cell survival (Solzak et al., 2013; Suizu et al., 2009), so increased levels of TTC3 could result in reduced cell numbers, as we observed in the frontal bone primordia of Dp1Tyb embryos. Finally, DSCAM is a cell adhesion molecule which has been proposed to act as a receptor for NETRIN-1, a laminin-related protein which inhibits osteoblast differentiation (Liu et al., 2009; Sato et al., 2017), thus increased dosage of DSCAM could affect osteoblastogenesis.

The craniofacial phenotype of Dp1Tyb mice is characterised by a shortening of the snout, contraction of the palate, micrognathia, doming of the skull and overall reduction in size (Toussaint et al., 2021). This dysmorphology remains largely consistent in models that break down the initial Dp1Tyb region, though the severity of the phenotype diminishes with decreasing size of duplication. The exception to this is the Dp6Tyb mouse strain which showed a largely opposite phenotype with a dolichocephalic skull, i.e., longer and flatter than WT controls. Interestingly, combining Dp5Tyb and Dp6Tyb increased dosage resulted in Dp1Tyb-like brachycephaly greater than that in Dp5Tyb alone, showing that increased dosage of the gene(s) causing the Dp6Tyb phenotype interact in a non-additive way with genes in the other regions.

One of the key characteristics of the craniofacial dysmorphology in Dp1Tyb mice is midfacial hypoplasia, as observed by the contraction of the snout and palate. This cranial phenotype correlated with aberrant synchondroses in Dp1Tyb mice, and other strains in the mapping panel, suggesting that these phenotypes may be linked. We found that both ISS and SOS show premature fusion in Dp1Tyb mice as early as P21, with evidence of mineralisation, which is normally absent in WT C57BL6/J mice at this age (Vora et al., 2016). Notably, premature fusion of synchondroses has also been reported in humans with DS at 7 months of age (Benda, 1940). Midfacial hypoplasia is commonly observed in other syndromes such as Pfeiffer’s and Crouzon’s, and again strongly correlates with premature fusion of the synchondroses (Goldstein et al., 2014; Paliga et al., 2014; Tahiri et al., 2014). Furthermore, surgical immobilisation of the SOS in rabbits prevented growth of the skull along the anteroposterior axis (Rosenberg et al., 1997). This linkage between midfacial hypoplasia and premature fusion of synchondroses has led to two alternative hypotheses. Firstly, premature fusion of the midline synchondroses may prevent anteroposterior growth causing midfacial hypoplasia, brachycephaly and reduction in overall cranial size (Paliga et al., 2014; Tahiri et al., 2014). Alternatively, brachycephaly and midfacial hypoplasia may place the opposing edges of the synchondroses close together, increasing local tissue stiffness and thereby causing premature ossification of the structure (Mauney et al., 2004; Sathi et al., 2015).

The key structures affected by the dysmorphic phenotype of Dp1Tyb mice from E13.5 to 16 weeks of age, are bones and structures of NC origin, such as the facial bones, frontal bones and the ISS (McBratney-Owen et al., 2008; Wu et al., 2017). We found that Dp1Tyb frontal bone primordia were reduced in size and cellularity at E13.5 and that the NC-derived cells of the frontal prominence which will give rise to these primordia, had reduced proliferation at E9.5. All these phenotypes were dependent on three copies of *Dyrk1a*. We hypothesise that increased dosage of *Dyrk1a* leading to reduced proliferation of NC-derived cells causes reduced size of the facial and frontal bones, and aberrant fusion of the synchondroses, which together lead to brachycephaly and midfacial hypoplasia.

Earlier studies of the Ts65Dn mouse model of DS had postulated that craniofacial dysmorphology in these mice was also caused by increased dosage of *Dyrk1a* and a neural crest defect (McElyea et al., 2016; Roper et al., 2009). In one of these studies, explants of Ts65Dn neural crest cells showed defective migration (Roper et al., 2009), in contrast to our findings in Dp1Tyb mice. Ts65Dn mice have not only an extra copy of 128 Hsa21-orthologous genes from Mmu16, corresponding to all the genes in the Dp2Tyb and Dp3Tyb regions and of some of the genes in the Dp9Tyb region, but also have an extra copy of 46 genes from Mmu17 that are not orthologous to Hsa21 (Duchon et al., 2011; Reinholdt et al., 2011). It is therefore not possible to know if phenotypes in this strain are derived from increased dosage of Hsa21-orthologous genes, or the latter, non-orthologous ones, or a combination of both. This key genetic difference could account for our observation that neural crest migration was unaffected in Dp1Tyb embryos, in contrast to the results reported for Ts65Dn mice.

We observed that the frontal bone primordia of Dp1Tyb embryos expressed lower levels of *Runx2*, a key regulator of ossification (Komori et al., 1997; McGee-Lawrence et al., 2014). This deficit may also contribute to the craniofacial dysmorphology of Dp1Tyb mice. Since this reduction in *Runx2* expression does not depend on three copies of *Dyrk1a*, it may reflect a different pathological mechanism acting in parallel with DYRK1A-driven perturbations. Certainly, DYRK1A is not the whole story when it comes to the DS phenotype, and whether the other genes, which we have mapped to other regions, operate via DYRK1A-associated molecular pathways or more in parallel, interacting at the level of differentiation or morphogenesis, remains to be determined.

In conclusion, we have shown that craniofacial dysmorphology in the Dp1Tyb mouse model of DS, which resembles that seen in humans with DS, is caused by at least four genes, one of which is *Dyrk1a*. Furthermore, we show that increased dosage of *Dyrk1a* results in impaired proliferation of NC cells and subsequent reduced size and cellularity of frontal and facial bones, as well as abnormal mineralisation of midline synchondroses. Together, these perturbations give rise to the brachycephaly and midfacial hypoplasia that typifies the cranium in DS.

## Materials and Methods

### Mice

C57BL/6J.129P2-Dp(16Lipi-Zbtb21)1TybEmcf (Dp1Tyb), C57BL/6J.129P2-Dp(16Mis18a-Runx1)2TybEmcf (Dp2Tyb), C57BL/6J.129P2-Dp(16Mir802-Zbtb21)3TybEmcf (Dp3Tyb), C57BL/6J.129P2-Dp(16Mir802-Dscr3)4TybEmcf (Dp4Tyb), C57BL/6J.129P2-Dp(16Dyrk1a-B3galt5)5TybEmcf (Dp5Tyb), C57BL/6J.129P2-Dp(16Igsf5-Zbtb21)6TybEmcf (Dp6Tyb), C57BL/6J.129P2-Dp(16Lipi-Hunk)9TybEmcf (Dp9Tyb), and C57BL/6J.129S6-Dp(16Cbr1- Fam3b)1Rhr (Ts1Rhr) mice have been described before (Lana-Elola et al., 2016; Olson et al., 2004). Dp5Tyb and Dp6Tyb mice were inter-crossed to generate Dp5Tyb/Dp6Tyb double mutant mice. Dp1Tyb mice were crossed to *Dyrk1a*^tm1Mla^ mice (*Dyrk1a*^+/-^, *Dyrk1a*KO) (Fotaki et al., 2002) to generate Dp1Tyb/*Dyrk1a*^+/+/-^ (Dp1Tyb/*Dyrk1a*KO) mice. Since both Dp1Tyb and *Dyrk1a*KO mice are poor breeders, it was not possible to generate sufficient numbers of experimental mice with this cross. Thus, we identified mice in which a crossover had placed the *Dyrk1a*KO mutation in cis on the same chromosome as the Dp1Tyb mutation. The resulting Dp1Tyb/*Dyrk1a*KO mice were bred against WT mice generating double mutants and WT animals in Mendelian ratios (1:1). Dp3Tyb mice were crossed to *Dyrk1a*KO mice to generate Dp3Tyb/*Dyrk1a*^+/+/-^ (Dp3Tyb/*Dyrk1a*KO) mice. All mice were bred at the MRC Harwell Institute except Dp3Tyb/*Dyrk1a*KO mice which were bred at The Francis Crick Institute. All mice were backcrossed to C57BL/6J for at least 10 generations. All animal work was approved by the Ethical Review panel of the Francis Crick Institute and was carried out under Project Licences granted by the UK Home Office. Numbers of protein-coding genes in different mouse strains were determined from mouse genome assembly GRCm39.

### Embryo collection

The day a vaginal plug was found was termed embryonic day 0.5 (E.5). Embryos were collected at E8.5, E9.5, E13.5, E16.5 and E18.5. Embryos were dissected out and those younger than E14.5 were immediately placed into ice-cold PBS. Embryos older than E14.5 were decapitated. For Haematoxylin and Eosin (H&E) staining and X-ray μCT analysis, heads were fixed in 4% PFA at 4°C overnight. For RNAscope analysis, heads were fixed in 10% Neutral buffered formalin (NBF) for 16-32 h at room temperature (RT). For immunofluorescence assays whole embryos at E9.5 were fixed for 1 h at RT.

### X-ray microcomputed tomography

For analysis of post-natal mice (16 weeks of age or P21), we used 10 mutant animals of each strain (5 female, 5 male) and 10 WT control age- and sex-matched mice from the same litters. Mice were killed via cervical dislocation. Heads were dissected from bodies and fixed for 24 h in 4% Paraformaldehyde (PFA). Heads of 16-week old and P21 mice were rinsed in PBS for 2 x 1 h and then scanned in a SCANCO μCT 50 at a 20 μm resolution, at 70 kV and with 500 3 s projections. Heads of embryonic mice were rinsed in PBS for 2 x 1 h and then scanned in a SCANCO μCT 50 at a 9 μm resolution, at 90kV and with 1500 3 s projections.

### Landmark-based morphometric analysis

68 three-dimensional landmarks for the crania and 17 landmarks for the mandible were used as previously defined (Hallgrímsson et al., 2007; Toussaint et al., 2021). Landmarks were place manually on a 3D volumetric reconstruction of the μCT images using the Microview (Parallax innovations) software; the same operator placed landmarks for all strains and was always blinded to genotype. Positioning of landmarks was checked using orthogonal planar views of the scanned subject. The same landmark sets were used for all mice. Principal component analysis (PCA) was conducted to visualise group shape separation. Statistical significance of shape differences was quantified by using the Procrustes Distance Multiple Permutations test at 1000 iterations. Normalised centroid size, defined as the square root of the sum of the squared distances of all landmarks from the centroid (the average x, y, z co-ordinate for each landmark dataset), divided by the number of landmarks was calculated using MorphoJ (Klingenberg, 2011). Statistical significance of size differences was calculated using a 2-tailed unpaired t-test.

### Landmark-free morphometric analysis

Landmark free analysis of P21, E18.5 and E16.5 mice was conducted as previously described (Toussaint et al., 2021). In brief, thresholding was applied to the µCT images extracting and generating a binary mask of the skull structures. Mandibles were separated from the rest of the cranium using segmentation based on bone density. Meshes of both the mandible and cranium were generated and aligned using Procrustes-based superimposition. Aligned meshes were subjected to atlasing using Deformetrica. Finally, outputs of atlas construction were analysed through PCA and a stratified k-fold cross validation test performed on the output PCA data. Significance of the classification score was then tested using a multiple permutations test at 1000 iterations.

### Paraffin embedding

Following fixation, embryos were washed in PBS for 3 x 30 min, then dehydrated through a series of ethanol solutions of 30%, 50%, 70%, 80%, 90%, and 100% for 1 h each at RT on a nutator and subsequently placed in cassettes. Using a Leica ASP300 tissue processor, embryos were place in xylene for 3 x 1 h and Ultraplast Wax for 3 x 1 h at 63°C. Following this, heads were placed in metal moulds in the desired orientation, embedded in paraffin wax and left to cool on an ice block. Once the wax had solidified, they were stored at 4°C until required.

### Microtome sectioning

Embryos embedded in paraffin blocks were trimmed down to an appropriate size using a razor blade and then sectioned on a Leica RM2245 microtome with disposable Leica microtome blades at a 10 μm thickness. Serial sections were placed into a 42°C deionised water bath to help with mounting onto Superfrost Plus slides. Slides were dried overnight on a heat block. For RNAscope, sections were cut to 5 μm and dried overnight at RT.

### Histology

10 μm sections were obtained and stained with haematoxylin and eosin for visualisation of the frontal bone mesenchymal condensation. For immunofluorescence, antigen retrieval was conducted by incubating whole E9.5 mouse embryos in Tris-EDTA buffer (10 mM Tris Base, 1 mM EDTA, 0.05% Tween 20, pH 8.0) for 1 h. The embryos were washed in PBS and underwent permeabilisation in 0.5% PBST for 30 min. They were washed again and subsequently blocked with 20% normal goat serum (NGS) for 1 h and incubated overnight at 4°C with rabbit polyclonal antibody against Sox10 (Proteintech, 10422-1-AP) in 20% NGS. Sections were washed with PBS, blocked in 20% NGS for 1 h and incubated with goat anti-rabbit IgG Alexa488 and DAPI overnight at 4°C. Finally, sections were washed with PBS and mounted with ProLong Diamond Antifade Mountant (ThermoFisher).

### Neural crest cell migration assay

Primary mouse cranial neural crest explant cultures were performed as previously described (Gonzalez Malagon et al., 2019). Embryos were harvested at E8.5 and immediately placed into ice-cold PBS. The anterior neural plate border region was dissected out and plated onto a glass-bottomed, 24-well tissue culture plate (Ibidi) coated with 1 µg/ml fibronectin. Explants were cultured overnight at 37°C and 5% CO_2_ in neural crest media containing Dulbecco’s modified Eagle’s medium, 15% fetal bovine serum, 0.1 mM minimum essential medium nonessential amino acids, 1 mM sodium pyruvate, 55 µM β-mercaptoethanol, 100 units/ml penicillin, 100 units/ml streptomycin and 2 mM L-Glutamine, conditioned by growth-inhibited STO feeder cells. Phase contrast imaging was carried out using an IX 81 microscope (Olympus), with a Cascade II 512B camera (Photometrics), and a 10x UPlanFL NA1.45 objective lens for 18 h every 5 min. Cells were tracked manually by following their nuclear position using the Manual Tracking plugin (Fiji). Cell tracks were imported into Wolfram Mathematica to analyse mean track speed and mean track persistence using the Chemotaxis Analysis Notebook v1.5β (G. Dunn, King’s College London, England, UK). Mean track speed is the average cell speed over the whole track length. Persistence was defined as the ratio of the cell’s total displacement to the total distance travelled over a time interval of 20 min. Mean track persistence is the average persistence over multiple time intervals (Law et al., 2013). The metrics were analysed using Prism 8 (GraphPad) to generate graphs and to calculate statistical significance using Tukey’s multiple comparisons test.

### RNAscope *in situ* hybridisation

Multiplex RNAscope *in situ* hybridisation was conducted using the RNAscope multiplex V2 reagent kit (ACD) and probes against *Runx2* (414021), *Osterix* (403401-C2) and *Msx2* (421851, all from ACD). Slides were imaged as a tile scan using a Vectra Polaris Automated Quantitative Pathology Imaging System (Akoya Biosciences) at 20x magnification. Regions of interest were selected using the whole slide scan viewer Phenochart (Akoya Biosciences) for unmixing. Tiles of the image were spectrally unmixed using inForm Tissue Analysis Software (Akoya Biosciences), stitched together in Qupath, and the frontal bone mesenchymal condensation was outlined. The total area of fluorescence for each gene in this area was calculated using the cell detection tool and divided by the total area to give expression per μm^2^. The average measurements for each slide were analysed using one-way ANOVA with Tukey’s multiple comparisons to calculate statistical significance.

## Acknowledgements

We thank Heather Cater and Sara Wells of the Mary Lyon Centre for breeding mice. We thank Alasdair Edgar for assistance with histology.

## Author contributions

Conceptualization: VLJT, JBAG

Formal analysis: YR

Funding acquisition: VLJT, JBAG, EMCF

Investigation: YR, DG, LD

Project administration: VLJT, JBAG

Resources: EL-E, SW-S, MK

Supervision: VLJT, JBAG, KJL

Visualisation: YR, VLJT, JBAG

Writing - original draft: YR, VLJT, JBAG

Writing - review & editing: YR, LD, KJL, VLJT, JBAG

## Competing interests

The authors declare no competing interests.

## Funding

VLJT and EMCF were supported by the Wellcome Trust (grants 098327 and 098328) and VLJT by the Francis Crick Institute which receives its core funding from Cancer Research UK (FC001194), the UK Medical Research Council (FC001194), and the Wellcome Trust (FC001194). YR was supported by a PhD studentship from the Francis Crick Institute and King’s College London. JBAG was supported by King’s College London. KJL and MK were supported by BBSRC (BB/R015953/1). LD was supported by a Medical Research Council Doctoral Training Programme studentship. For the purpose of Open Access, the authors have applied a CC BY public copyright licence to any Author Accepted Manuscript version arising from this submission.

## Supplementary Data

**Table S1.**
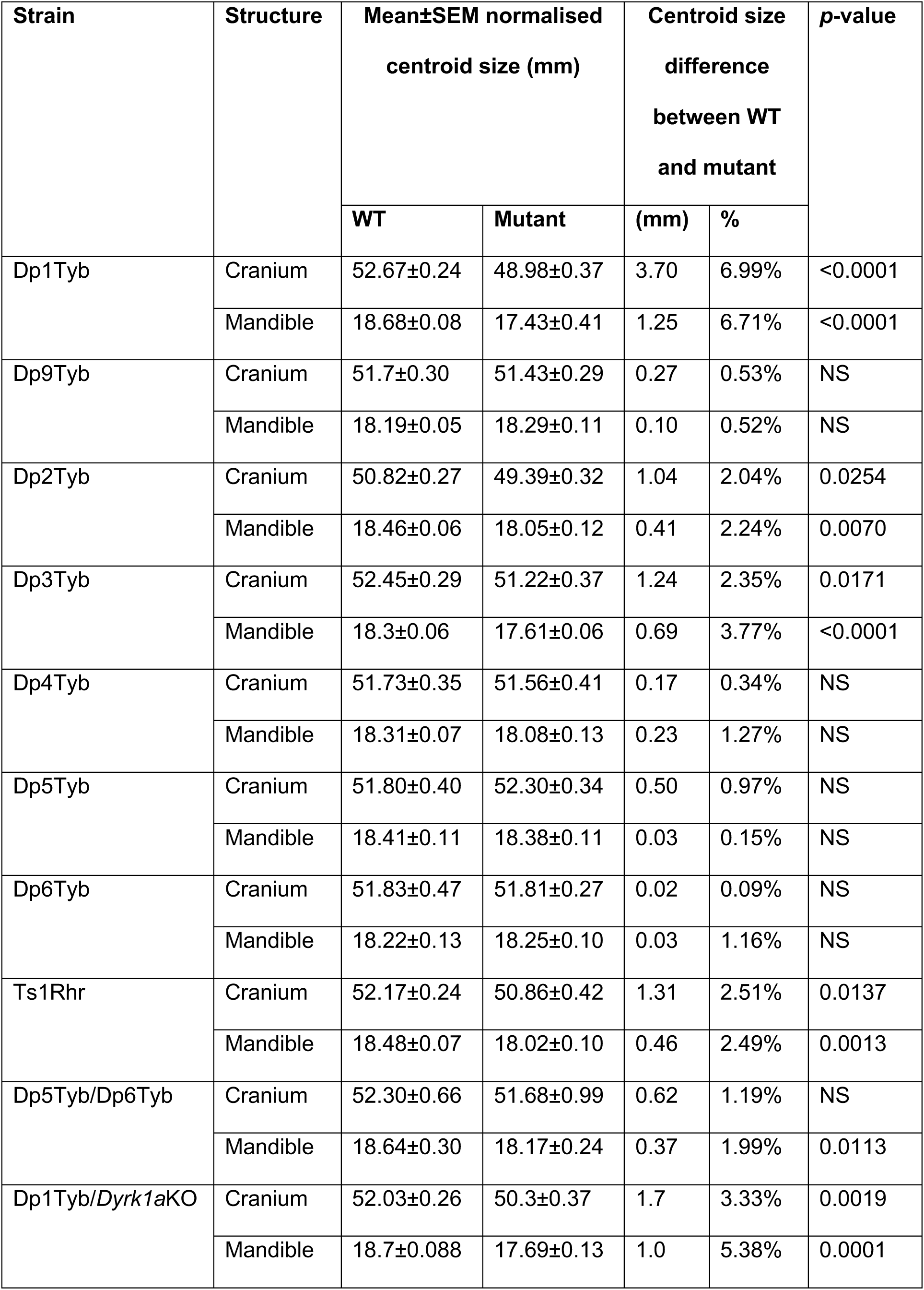
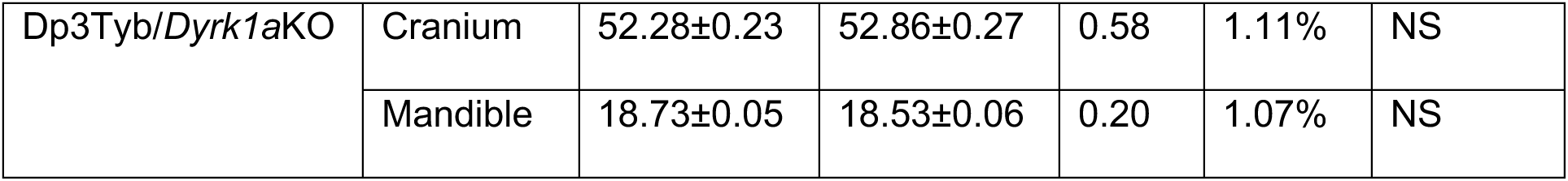
Centroid sizes of crania and mandibles from all mouse strains analysed in the genetic mapping panel. Comparison of centroid sizes of crania and mandibles in 16-week old mice from all strains used in the genetic mapping study and their respective WT controls. NS, not significant (*p*-value > 0.05).

**Figure S1.**
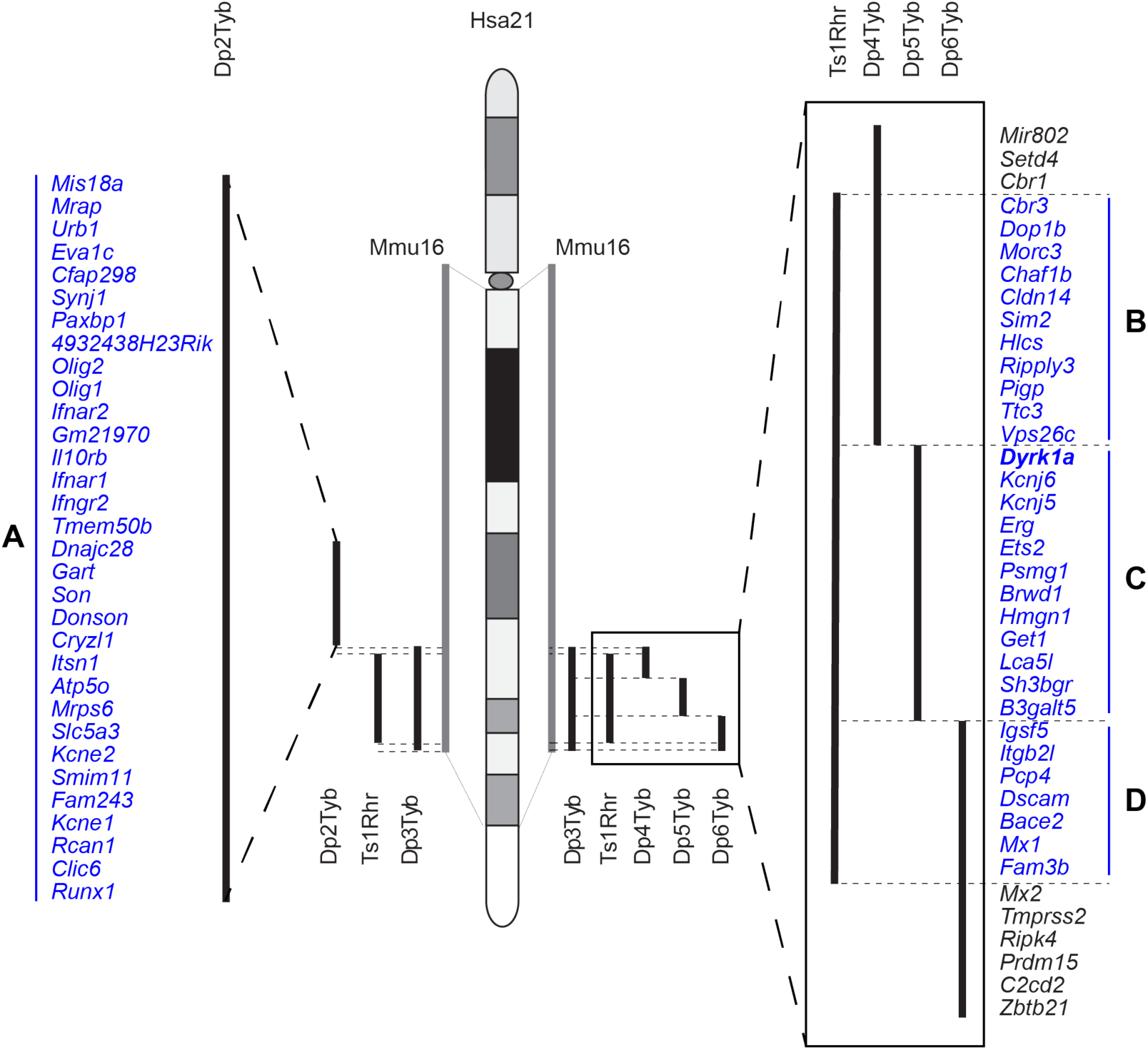
Regions of Hsa21 containing dosage-sensitive genes causing craniofacial dysmorphology. Diagram of Hsa21 showing main cytogenic regions (rectangles of different shades) and the centromere (oval). Grey bars indicate the Hsa21-orthologous region of Mmu16. Black lines on the right hand of the diagram indicate regions of Mmu16 duplicated in the Dp3Tyb, Ts1Rhr, Dp4Tyb, Dp5Tyb and Dp6Tyb mouse strains. Black lines on the left hand of the diagram indicate regions of Mmu16 duplicated in the Dp3Tyb, Ts1Rhr and Dp2Tyb strains. These regions are expanded showing all known protein coding genes within them and one microRNA gene (*Mir802*). Genes in blue lie within the four regions shown in this study to cause craniofacial phenotypes when present in three copies: regions A (*Mis18a* - *Runx1*), B (*Cbr3* - *Vps26c*), C (*Dyrk1a* - *B3galt5*) and D (*Igsf5* - *Fam3b*). The *Dyrk1a* gene (bold) is required in three copies for the full phenotype. Genes not within regions that cause craniofacial dysmorphology are in black.

## Supplementary Videos

**Video S1**

Cranial morph between WT and Dp1Tyb skulls at P21 with size regressed out.

**Video S2**

Cranial morph between WT and Dp1Tyb skulls at E18.5 with size regressed out.

## Notes

### Competing Interest Statement

The authors have declared no competing interest.

